# Migration patterns and hybridization within the Asian stonechat complex in response to a major geographical barrier

**DOI:** 10.1101/2025.10.15.682589

**Authors:** Tianhao Zhao, Yury Anisimov, Wieland Heim, Gang Song, Valentina Anisimova, Nyambayar Batbayar, Christen M. Bossu, Andrea Bours, Siqiao Chen, Wenjia Chen, Batmunkh Davaasuren, Dule, Shengjing Jiao, Xiaolu Jiao, Magnus Hellström, Georg Langebrake, Zhuo Li, Shu-Yueh Liao, Aitao Liu, Zongzhuang Liu, Jacob Roved, Xinyuan Wang, Matthias H. Weissensteiner, Guannan Wen, Dezhi Zhang, Guoming Zhang, Yifang Zhang, Kristen Ruegg, Miriam Liedvogel, Staffan Bensch, Bregje Wertheim, Fumin Lei, Barbara Helm

## Abstract

Long-distance avian migration is thought to be under strong natural selection. Facing geographical barriers, migrants display various patterns considered to be adaptive. For example, they may detour along either side around the barrier or cross it, requiring specialized behavioral adaptations. Variations within closely related taxa are excellent sources for understanding the evolutionary background of migration and how barriers are shaping migration routes.

In Asia, some species are assumed to have a migratory divide in response to the major geographical barrier, the Qinghai-Tibet Plateau (QTP), including the stonechat taxa (Siberian Stonechat *Saxicola maurus maurus* and Amur Stonechat *S*. *stejnegeri*). As they detour along either side of the QTP, these taxa are believed to disfavor a crossing over the highland. However, the more southernly distributed Tibetan Stonechat (*S. m. przewalskii*) breeds on the QTP, suggesting adaptation to high elevation. To investigate migration patterns and the potentially associated genetic differences, we studied migration routes and population genetics of four populations around the assumed migratory divide in Russia and Mongolia, and of one from the QTP in China. Our results confirmed the existence of a migratory divide between *maurus* and *stejnegeri*, albeit with extensive hybridization. We observed both the hypothesized western and eastern routes, but also found individuals employing intermediate routes crossing the QTP, of which two-thirds were clear hybrids. Meanwhile, *przewalskii* followed a highland-crossing route and was genetically differentiated from *maurus* and *stejnegeri*.

The diverse migration routes among Asian stonechats show differential responses towards the geographical barrier. The intermediate route may be associated with hybridization, and its conditional viability may facilitate gene flow between *maurus* and *stejnegeri*.

The Asian stonechat complex thus offers great opportunities for novel research of the genetics and evolution of migration. The specific evolutionary background associated with inhabiting and crossing the QTP can offer new perspectives in this field.

**Teaser text:** Migratory divides can arise in birds because alternative routes around migratory barriers would select for behaviors to restrict hybridization. Hybrids of parental types that employ alternative routes are hypothesized to embark on intermediate routes that would expose them to suboptimal conditions, resulting in post-zygotic reproductive isolation. However, this hypothesis is challenged when a sister taxon actually breeds on the geographical barrier. This is the case in the Asian stonechat complex that breeds near or on the Qinghai-Tibet Plateau (QTP), the roof of the world. We demonstrated a migratory divide in central Siberia to Mongolia for race *maurus* and *stejnegeri*, yet showed also evidence for extensive hybridization. Hybrids migrated along a newly discovered intermediate route, seemingly viable and overlaps with the migration trajectory of race *przewalskii* over the eastern part of the QTP.

The Asian migratory divide relative to the QTP thus provided new insights to the evolution of landbird migration.

## Introduction

Migration behavior exhibits a wide diversity across avian taxa, encompassing its occurrence, timing, direction, routes, and distances. Our accumulated knowledge on migration patterns has revealed substantial inter- and intra-specific variation, which is often associated with genetic differences and evolutionary histories (Liedvogel et al., 2011; Ruegg et al., 2017; Justen & Delmore, 2022).

Migratory divides, i.e. areas where two closely related taxa that exhibit different migration routes meet, mate and likely also hybridize, are focal points of spatial diversity during migration. Such migratory divides have been documented in birds of all major global flyways, e.g., in the European-African Flyway within Willow Warblers (*Phylloscopus trochilus*) and Eurasian Blackcaps (*Sylvia atricapilla*) (Bensch et al., 1999, 2009; Caballero-López et al., 2021; Sokolovskis et al., 2023; Delmore et al., 2020a; A. Helbig, 1996; A. J. Helbig et al., 1994); in the Americas Flyway within Swainson’s Thrush (*Catharus ustulatus*) (Ruegg & Smith, 2002); in the Atlantic Flyway within Red-necked Phalarope (*Phalaropus lobatus*) (van Bemmelen et al., 2019); and in the Central Asian Flyway within the Barn Swallow (*Hirundo rustica*) (Scordato et al., 2020; Turbek et al., 2022). There is also a presumed east-west Siberian migratory divide in central Siberia described in, for example, the Siberian/Amur Stonechats (*Saxicola maurus, S. stejnegeri*), Citrine Wagtail (*Motacilla citreola*), and Siberian Bluethroat (*Luscinia svecica*) (Irwin & Irwin, 2005; Strubbe et al., 2025).

It has been frequently proposed that phenotypic variation in migration behaviors, especially within migratory divides, is associated with geographical barriers, for example, the Sahara Desert, the Mediterranean Sea, the Rocky Mountains, and the Qinghai-Tibet Plateau (Irwin & Irwin, 2005; Justen et al., 2024; Sokolovskis et al., 2023; Turbek et al., 2022; Zhao et al., 2020, 2024). The challenges from either the lack of opportunities to rest, low oxygen, bad weather, or the lack of refueling sites, result in a higher fatality rate when crossing directly over geographical barriers than using longer detours to bypass (Bishop et al., 2015; Irwin & Irwin, 2005; Sjöberg et al., 2021). However, some avian taxa can migrate directly across such barriers (Hawkes et al., 2011; Lindström et al., 2021; Sjöberg et al., 2021; Yu et al., 2024).

The migration direction of largely solitary migrating songbirds is under partial genetic control and undergoes selection through adaptation processes (Justen & Delmore, 2022; Liedvogel et al., 2011). Since the first study on captive Eurasian blackcaps, it has been hypothesized that the direction of migration is controlled by a few genes (Helbig, 1991). Research on revealing the genetic basis of migration direction has been carried out in several songbird species that have a migratory divide (Bascon-Cardozo et al., 2022; Delmore et al., 2020; Bensch et al., 1999, 2009; Ruegg et al., 2002, 2014; Helbig, 1996; Lundberg et al., 2017; Sokolovskis et al., 2023; Turbek et al., 2022; Zhao et al., 2020). Interestingly, the genetic signals associated with phenotypic variations detected hitherto typically differ between study systems (Caballero-López & Bensch, 2024). Parallel evolution histories and differential environmental factors along different flyways may have promoted convergent evolution (Bensch et al., 2023; Caballero-López & Bensch, 2024).

Conspecific phenotypic variation in migration patterns can also facilitate the speciation process. Differences in migration direction can act as a barrier for gene flow. Helbig (1991) proposed that in Eurasian Blackcaps, crossbred offspring from parental individuals migrating to Southeast and Southwest in autumn would follow an intermediate route south across the Alps and the Mediterranean Sea, and this was hypothesized as suboptimal and associated with high fatality. This is hypothesized to function as a mechanism of post-zygotic reproductive isolation limiting the gene flow between populations (Helbig, 1991), but viable intermediate routes have since been demonstrated to exist without compromising survival in the wild (Delmore et al. 2020a, 2023). Alternatively, or in addition to possible differentiation due to spatial factors, temporal factors can also restrict gene flow (Bearhop et al., 2005; Turbek et al., 2022). For example, allochrony in spring arrival phenology can be a pre-zygotic reproductive isolation mechanism, resulting in a trend of assortative mating between parents with similar arrival times (Bearhop et al., 2005; Turbek et al., 2022).

While major advances have been achieved in studying the evolution and adaptation of migration traits in the recent years, Asian landbird species have rarely been taken into consideration (Irwin & Irwin, 2005; McKinnon & Love, 2018; Turbek et al., 2022). In Asia, the Qinghai-Tibet Plateau (QTP) and its surrounding mountains and deserts potentially present a significant barrier to migration in species breeding in Siberia, within the range of QTP, and wintering south of QTP (Zhao et al., 2024). With an average elevation over 4000 m above sea level (a.s.l.) and an area over 2.5 million km^2^, the massive plateau demands efficient respiration rate and usage of oxygen, tolerance of changeable weather, and endurance of crossing large barren areas with restricted refueling opportunities (Hawkes et al., 2011; Kumar et al., 2020). Species that are migrating across it at high altitude, e.g., the Bar-headed Goose (*Anser indicus*), the Demoiselle Crane (*Grus virgo*), and the Brown-headed Gull (*Chroicocephalus brunnicephalus*), exhibit special behavioral and physiological adaptations (Bishop et al., 2015; Galtbalt et al., 2022; Mi et al., 2022; Yu et al., 2024; Prins & Namgail, 2017). Although rarely described and discussed, several small highland-specialized songbirds are believed to employ a migration route across the QTP: White-throated Bushchat (*Saxicola insignis*) occasionally occurs on the QTP during the spring migration season (Urquhart, 2010; Collar 2020); Siberian stonechats and Rosy Pipits (*Anthus roseatus*) have been documented to fly across the Himalayas down to Bhutan (Proud 1949; Spierenburg, 2005). Otherwise, the majority of Siberian breeding passerine species and those breeding on QTP migrate along detoured routes either along the eastern or western side of the QTP (Irwin & Irwin, 2005; Kumar et al., 2020; Zhao et al., 2024).

Here, we introduce the common stonechat complex (*Saxicola* sp.) in Asia as a novel study system for studying phenotypes and genetic correlations of migratory birds facing the QTP barrier. The common stonechat complex occurs in Africa, Europe and Asia, with an unresolved taxonomy status (Drovetski et al., 2004; Illera et al., 2008; van Doren et al., 2017; Zink et al., 2009), including the Asian clade (Opaev et al., 2018). To simplify the taxonomical reference in this manuscript, we will use *maurus*, *stejnegeri*, and *przewalskii* to refer to each of the stonechat taxa.

Regarding the migration phenotype, there appears to be high diversity among Asian stonechats (**Figure 1**). The nominate subspecies of Siberian Stonechat *maurus* and Amur Stonechats *stejnegeri* potentially share a migratory divide in central Siberia, ranging south down to Lake Baikal and potentially further into Mongolia (Irwin & Irwin, 2005). A tracking study of the Hokkaido (Japan) population of Amur Stonechat revealed that they migrate through eastern China (Yamamura et al., 2017, 2025). Moreover, the Tibetan Stonechat *przewalskii* breeding on the QTP and central to southwest Chinese mountain regions are believed to be migratory, but their migration patterns remain unclear (Urquhart, 2010). In contrast, the Indian Stonechat (*S*. *m*. *indicus*) breeding in northeast India to Pakistan are believed to be sedentary (Urquhart, 2010). The ambiguous differentiation on color morphs between Siberian and Amur stonechat (Hellström & Norevik, 2014) further lead to low accuracy in identifying them during the non-breeding season, and thus, the migration routes for either of the species or subspecies remain unclear. Regarding genetic differentiation, previous studies on the stonechat taxa in Asia mainly conducted phylogenetic analyses on mitochondrial DNA (Illera et al., 2008; Opaev et al., 2018; Zink et al., 2009), and did not associate migration routes with population genetic structural differences (Justen et al., 2022; van Doren et al., 2017).

**Figure 1.**
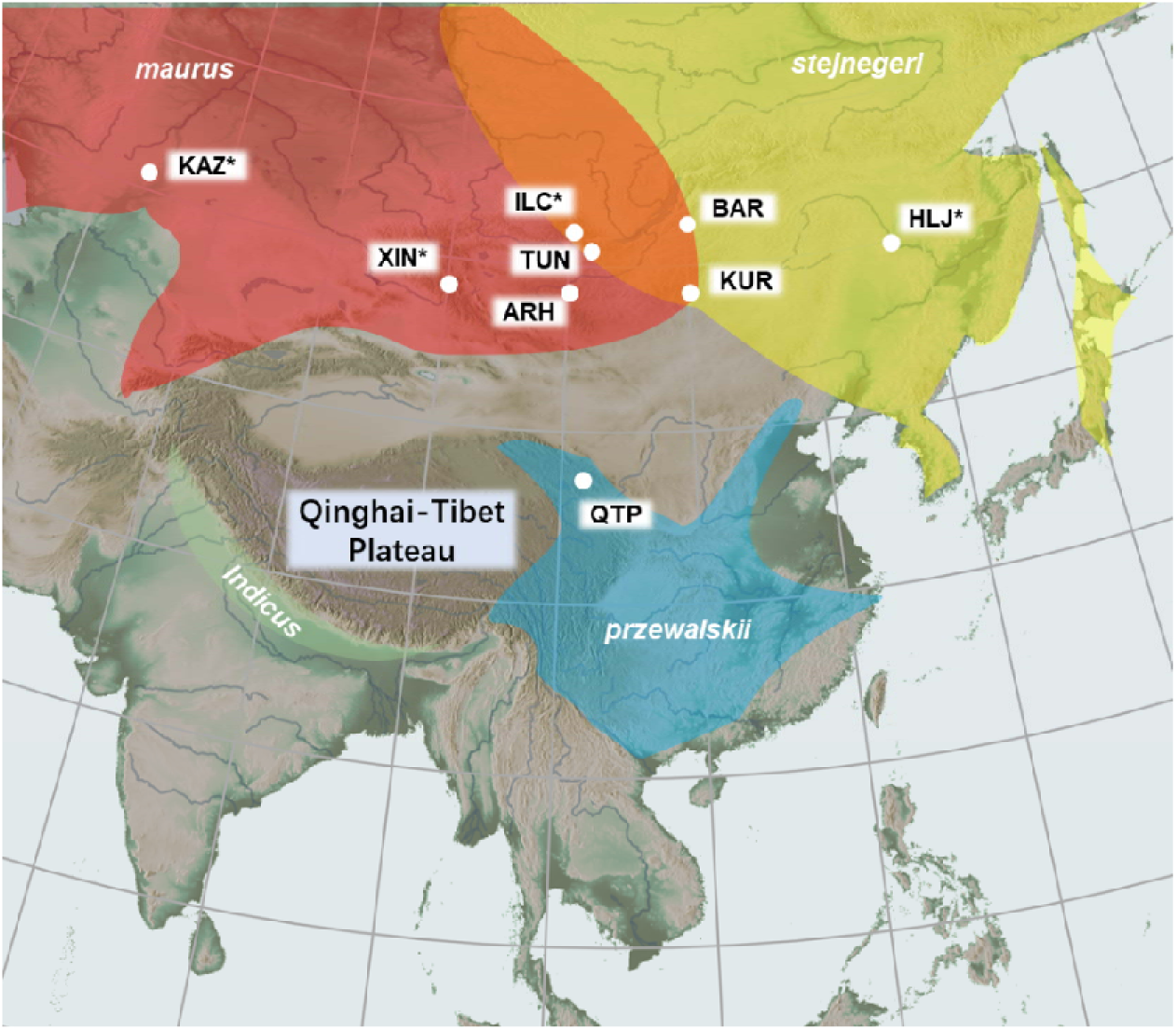
Schematic distribution map of stonechat taxa breeding in Asia (with a combined reference to citizen science data from eBird.org and www.birdreport.cn, Opaev et al., 2018, and IUCN redlist distribution data). Four taxa are displayed on the map: *maurus* (red), *stejnegeri* (yellow), *przewalskii* (blue), and *indicus* (green); the sedentary *indicus* race was not included in this study. The overlapped area between *maurus* and *stejnegeri* represent the location of the hypothesized contact zone between the two taxa. The white dots represent the sampling sites involved in this study: Kazakhstan (KAZ), Xinjiang, China (XIN), Ilchir lake, Russia (ILC); Tunka valley, Russia (TUN), Barguzin valley, Russia (BAR), Arhangai valley, Mongolia (ARH), Khurkh, Mongolia (KUR), Qinghai, China (QTP), Heilongjiang, China (HLJ). Location names with asterisks indicate that only genomic data was collected at these sites, and no individual migration tracking with geolocators was conducted. In some contexts, capitalized names (e.g., TUN) will be used as short form; locations with full name (e.g., Tunka) will be used in geolocator data reports; names of the taxa (e.g., *maurus*) will be used in taxonomical discussions.

Taking advantage of the diverse migration behaviors of stonechats in Asia, we investigated the phenotypes and genotypes of migrants faced with the QTP. We focused on the two taxa breeding north of the QTP near the migratory divide (*maurus* and *stejnegeri*), and compared them to a third taxon breeding on the QTP (*przewalskii*). For genomic analyses, we added samples from additional populations (**Figure 1**). Our objectives are: 1. To document migratory phenotypic diversity using individual tracking methods; 2. To relate the migratory phenotypic variation to genetic diversity by employing whole genome re-sequencing. We aim to test the following hypotheses: i) There is a migratory divide between *maurus* and *stejnegeri* in its documented contact zone in Siberia, and possibly in Mongolia, and migrants from either side will detour to avoid crossing the QTP; ii) hybrids are rare, and if detected, employ an intermediate route compared to their parental phenotype; iii) The highland breeding population *przewalskii* will also avoid crossing the QTP during migration; iv) There is clear genomic differentiation among *maurus*, *stejnegeri* and *przewalskii*, based on an extended range of samples and in association with their different migration patterns.

## Methods

### Fieldwork

Fieldwork was carried out at six different sites, five of which (excluding Ilchir) included individual tracking of adult birds (**Figure 1**). For the contact zone between *maurus* and *stejnegeri* we selected three field sites west of the divide, i.e. in the Tunka Valley, Russia (102.33 – 102. 39E, 51.74 – 51.81N, 800m a.s.l.), at Ilchir Lake, Russia (100.9E, 51.9N, 1900m a.s.l.) and a valley located in the Arhangai Province, Mongolia (100.86 – 100.91E, 48.72 – 48.74N, 1900m a.s.l.); and two sites east of the divide, i.e. the Barguzin Valley, Russia (109.79 – 109.82E, 53.56 – 53.58N, 500m a.s.l.) and at Khurkh Valley, Mongolia (110.10 – 110.21E, 48.43 – 48.46N, 1400m a.s.l.). For *przewalskii*, the field site was in Menyuan Hui Autonomous County in Qinghai, China (101.30 – 101.54E, 36.64 – 37.43N, 3300m a.s.l.).

We conducted our fieldwork from 2020 to 2024 between late May and early July during the breeding season of the local stonechat populations. We first scouted the area to identify nest sites, and then applied a combination of mist nets, perch traps and ground bait traps near the nests to catch adult breeders. We equipped each individual with one type of light-level geolocators (specifics are introduced in the next section), and banded the individual with an aluminum ring and an individual-specific combination of color rings. The aluminum rings were provided by the National Bird Banding Centre of China, the Wildlife Science and Conservation Center of Mongolia, and the Bird Ringing Centre of Russia. After the handling, we released the birds back to their territories; the same individuals were spotted and captured by the same method to retrieve the loggers in later years if they returned. We also sampled blood from the brachial vein under the left wing for most of the captured individuals and stored it in 70% Ethanol for DNA extraction.

### Geolocator deployment and analysis

We used light-level geolocators provided by two manufacturers: The Intigeo-P30Z11-DIP-NOT light level recorder from Migrate Technology Ltd. (MT, 0.36 g (excluding harness), 14 x 6 x 3 mm, 11 mm light stalk); and the SOI-GDL2 logger from Swiss Ornithological Institute (SOI, 0.68 g (excluding harness), 8 x 6 x 4 mm 7 mm light stalk). Both logger types were programmed to store light levels every 5 min from July 15^th^ in the deployment year for as long as the battery lasted.

We deployed 115 geolocators (60 MT, 55 SOI) on 58 males and 57 females at five field sites between 2021 and 2023, and successfully retrieved 17 devices in subsequent seasons. Of those, 16 included complete data, and one from Arhangai, Mongolia, stopped working before the autumn migration (**Table 1**). The SOI logger data were extracted by the manufacturer; the MT logger data were extracted using the IntigeoIF interface and software (Migrate Technology Ltd.). We conducted the analysis in R 4.3.2 (R Core Team, 2023), following the manual for geolocator analysis (Lisovski et al., 2020). We used the SGAT package to analyze movement trajectories following the instructions on https://geolocationmanual.vogelwarte.ch/SGAT.html (Lisovski et al., 2020). Analysis details can be found in **Supplementary methods** and **Table S1**.

**Table 1.** Light-level geolocator deployment and re-capture from different field sites.

| Country | Location | deployment year | Logger type | deployment number |  | recollection number |  |
| --- | --- | --- | --- | --- | --- | --- | --- |
|  |  |  |  | male | female | male | female |
| Russia | Tunka (TUN) | 2021 | MT<sup>a</sup> | 14 | 6 | 3 | 1 |
|  | Barguzin (BAR) | 2021 | MT | 10 | 10 | 1 | 1 |
| China | Menyuan (QTP) | 2021 | MT | 9 | 11 | 1 | 4<sup>c</sup> |
|  |  | 2022 | SOI<sup>b</sup> | 7 | 8 | 1 | 1 |
| Mongolia | Khurkh (KUR) | 2023 | SOI | 8 | 12 | 0 | 1 |
|  | Arhangai (ARH) | 2023 | SOI | 10 | 10 | 2 | 1+1<sup>d</sup> |

a. MT stands for Migrate Technology Ltd.
b SOI stands for Swiss Ornithological Institute
c. One logger (CA903) was deployed in 2021 and re-collected in 2023
d. One female lost its logger upon re-capture; the other female carried a logger (29EN) that stopped
recording data before the autumn migration

To supplement a visualization of the environmental effect, we extracted the Normalized Difference Vegetation Index (NDVI) score at each individual’s breeding geolocation. We extracted the NDVI from (https://ee-zhangcheng-data1.projects.earthengine.app/view/ndvi-time-series-explorer), which used the MODIS-061_MOD13Q1 dataset with 16 days interval and 250m resolution from 2000 to 2024. We downloaded the NDVI data for the 250m*250m grid where the breeding site was located for each studied individual during the year of tracking, and plotted them together with the individual’s migration schedule.

### Whole-genome sequencing

To investigate the population genetic structures among Asian migratory populations, especially in the contact zone and the Qinghai population, we selected 77 samples from 7 geographical breeding populations (**Figure 1, Table S2**). No genomic samples were yet available from the Mongolian study sites (ARH, KUR). To strengthen population genetic analyses, we complemented samples from our field sites with historical samples from Kazakhstan (Justen et al., 2022) and Xinjiang, China, presumably representing *maurus*, and from Heilongjiang, China, presumably representing *stejnegeri*. Both Xinjiang and Heilongjiang samples were specimen samples archived in the Institute of Zoology, Chinese Academy of Science. Sample sizes are as follows: Kazakhstan (KAZ, n = 9), Xinjiang, China (XIN, n = 4), Ilchir lake, Russia (ILC, n = 2), Tunka valley, Russia (TUN, n = 18), Barguzin valley (BAR, n = 17), Heilongjiang, China (HLJ, n =4), Qinghai, China (QTP, n =23) (**Table S2**). The samples from Qinghai, China were sampled in 2020 and 2021; the samples from 2021 were from most of the individuals tracked in 2021. Of the 16 birds whose tracking data are included, we had genomic information for 11 individuals (TUN_09, TUN_10, TUN_16, TUN_17, BAR_05, BAR_10, QTP_04, QTP_07, QTP_08, QTP_09, QTP_18, **Table S2**).

We extracted DNA with the Qiagen DNeasy Blood & Tissue Kits. The QTP samples were sequenced at a mean depth of 15X at Berry Genomics Inc., on an Illumina Novaseq instrument using a PCR method for pair-end reads of 150 bp. The remaining samples were sequenced at a mean depth of 15-20X at BGI Genomics Inc. for BGI DNBseq (WGS) using a PCR method for pair-end reads of 100 or 150 bp.

To identify single-nucleotide variants, WGS reads of both platforms were processed with the *variant-calling* pipeline, a custom-built *snakemake* workflow (gmanthey/variant-calling v1.0; https://doi.org/10.5281/zenodo.15969023; Mölder et al., 2021). Detailed descriptions can be found in **Supplementary methods**. We used the reference genome that was assembled by van Doren et al., 2017 (GCA_900205225.1 on Genebank).

To investigate the population genetic structures and explore the hybridization level at the contact zone between *maurus* and *stejnegeri*, we used the SNP data sets for several population genetic analyses. We further ran a filter process in Plink 1.07 (Purcell et al., 2007), based on: 1. MAF > 0.05; 2. SNPs in LD were filtered with a window size of 50 SNPs (advancing 10 SNPs at a time) and a r^2^ threshold of 0.2. We also excluded individuals KAZ_02 and KAZ_06 as they were found to be hand-raised from the same brood with KAZ_01 and KAZ_05 in captivity, respectively.

To visualize the population structure and identify hybrid individuals, we used the Admixture software (Alexander et al., 2009) to run admixture analyses from a range of K = 1–5, with a biological hypothesis of K = 3 (*maurus*, *stejnegeri* and *przewalskii*). We ran the analyses with 30 iterations, and picked the output with lowest CV error for each K from all iterations as the final outputs. We also ran a PCA analysis using the same SNP dataset using the distance-matrix function in Plink, extracting the PC1 and PC2 to visualize population structure. Additionally, to determine the type of the hybrids in the contact zone (Barguzin, Tunka and Ilchir) between *maurus* and *stejnegeri*, we used the triangular package (Wiens, 2025) and selected 647 diagnostic SNP loci (difference threshold at 0.9) using KAZ+XIN as P1 (*maurus*) and HLJ as P2 (*stejnegeri*) to illustrate the hybrid index and interclass heterozygosity pattern among individuals from the contact zone.

To investigate the level of genetic differentiation between populations, we calculated the weighted F_ST_ between each pair of geographical populations using the --fst function in Plink. To report the population heterozygosity level, we used the --het function in Plink, and calculated the observed heterozygosity = 1 – O(HOM)/N, where O(HOM) stands for the observed homozygotic SNP loci, and N stands for the total number of SNPs. We did not include the invariant sites for the calculations. For both calculations, we combined the populations of Kazakhstan and Xinjiang, China as one (KAZ+XIN), as they were both far from the contact zone and represented the pure *maurus* genetic background. We also combined the Ilchir and Tunka populations as (TUN+ILC), because the Ilchir population only had two individuals, and they were both located on the western side of the contact zone and not far from each other. All the visualization was conducted in R 4.3.3 (R Core Team, 2024).

## Results

### 1. Migration routes

We acquired 17 autumn and 16 spring migration tracks of 16 individuals (8 males and 8 females) (**Figure 2, Figure S1**). From the four individuals recaptured from the Tunka population, west of Lake Baikal, two individuals followed the western route in both autumn and spring, and wintered in north India; the other two individuals followed an intermediate route in both seasons, where one wintered in central India and the other wintered on the south side of the east-range of the Himalayas. From the two individuals recaptured from the Barguzin population east of Lake Baikal, one individual followed the eastern route in both seasons and wintered in the lowland region in central Myanmar; the other individual followed an intermediate route and wintered in central India. The two individuals recaptured from the western Mongolian Arhangai population both followed the western route both in autumn and spring, and wintered in north India. The one individual recaptured from the eastern Mongolian Khurkh population followed an intermediate route during autumn and the eastern route during spring, and wintered on the south side of the middle-range of the Himalayas. All seven individuals recaptured from the Qinghai population migrated across the QTP in autumn and spring, and wintered around the eastern range of the Himalayas. The data precision did not allow us to judge whether these individuals wintered on or off the QTP; however, 4 out of 7 individuals had one or several detected stopover(s) around the Assam Plain during spring, which is connected with the QTP through the Yarlung Zangbo Grand Canyon.

**Figure 2.**
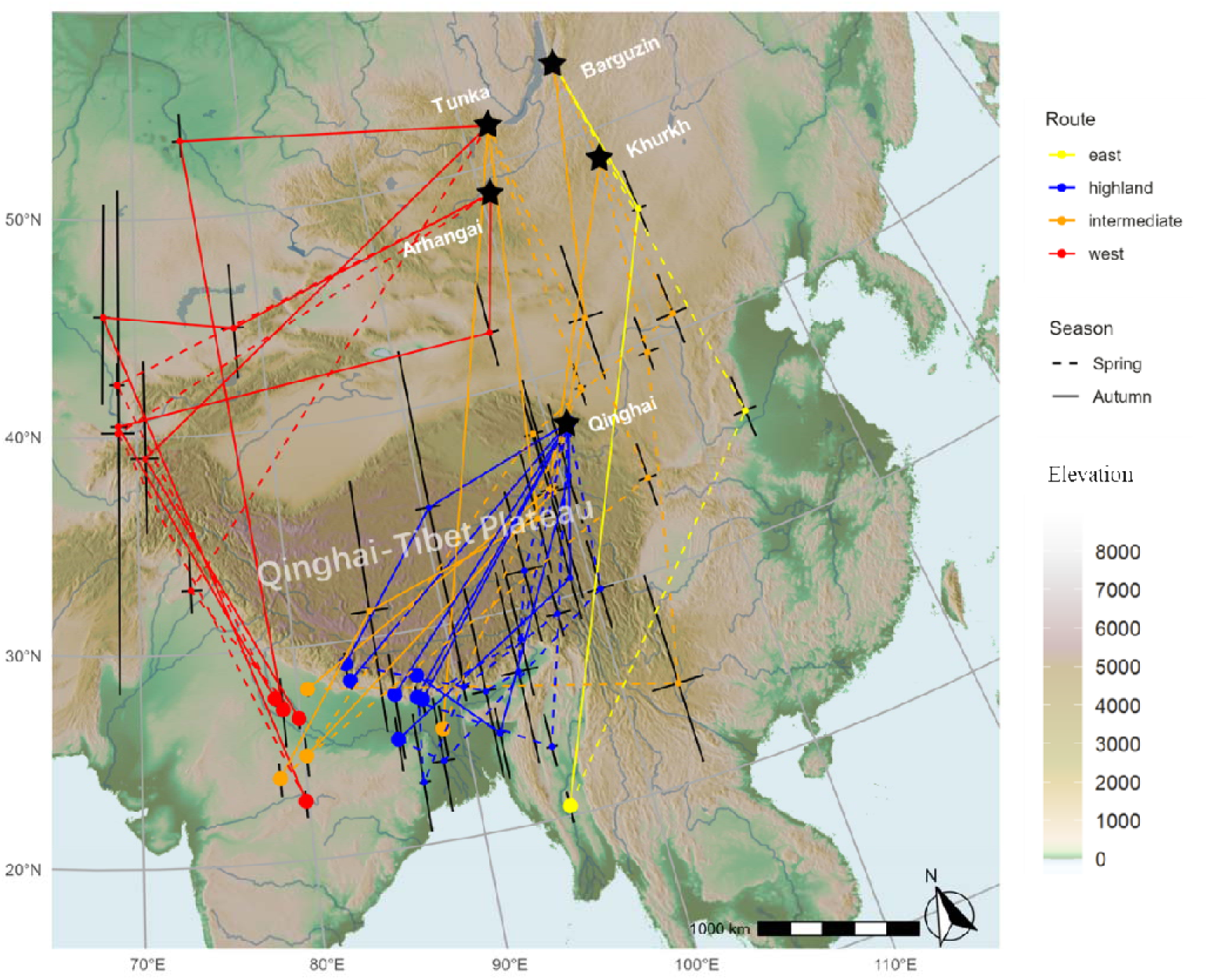
Migration routes of stonechats from different populations in Asia, estimated based on light-level geolocator data. The five deployment locations (breeding sites) are marked with names and black stars. The Qinghai-Tibet Plateau is labeled on the map. Four types of migration routes are subjectively classified and displayed in different colors. Wintering locations are displayed as large dots, whereas stopover sites are displayed as small dots. The center of the dot represented the median location estimated from the MCMC model in the SGAT analysis, whereas the error bars are drawn according to the 2.5% and 97.5% of the latitude/longitude estimation from the MCMC model. The color of the background map refers to the ground elevation. Potential autumn routes are drawn as solid lines, whereas potential spring routes are drawn as dashed lines. Routes are estimated as the shortest trajectory between detected stationary sites.

The migration distances of both seasons for each individual are shown in **Table 2**. The migration distance of the Tunka individuals was 1281±200 km (38%) longer than the bee-line distance, and of the Arhangai individuals, it was 1886±74 km (59%) longer. For the individual following the eastern route from Barguzin, their migration distance was 610 km (16%) longer than the bee-line distance. We calculated the autumn and spring migration distance separately for the individuals following an intermediate route, as their migration routes were not always consistent between seasons: in autumn, the migration distance was 409±133 km (11%) longer than the bee-line distance; and in spring, it was 966±477 km (27%) longer than the bee-line distance (**Table 2**, **Figure 2**, **Figure S2**)

**Table 2.** Basic information of each tracked individual and their migration spatio-temporal patterns.

| ID | Breeding location | Sex | Age at 1<sup>st</sup> capture | Route type | Year | Autumn departure | Autumn arrival | Autumn migration duration (day) | Spring departure | Spring arrival | Spring migration duration (day) | Bee-line distance (km) | Autumn migration distance (km) | Spring migration distance (km) | Prolonged autumn distance (km) | Prolonged spring distance (km) |
| --- | --- | --- | --- | --- | --- | --- | --- | --- | --- | --- | --- | --- | --- | --- | --- | --- |
| CA863 | Barguzin | F | 2 | East | 2021 | 2021-08-15 | 2021-10-01 | 47 | 2022-04-20 | 2022-05-24 | 34 | 3880 | 4082 | 4490 | 202 | 610 |
| 29AM | Khurkh | F | 2+ | Intermediate | 2023 | 2023-08-29 | 2023-09-29 | 31 | 2024-04-13 | 2024-05-04 | 21 | 3437 | 3692 | 5038 | 255 | 1601 |
| CA869 | Tunka | M | 3+ | Intermediate | 2021 | 2021-09-04 | 2021-10-12 | 38 | 2022-04-08 | 2022-05-06 | 28 | 3163 |  | 3461 |  | 298 |
| CA886 | Tunka | M | 2+ | Intermediate | 2021 | 2021-08-28 | 2021-09-23 | 26 | 2022-04-09 | 2022-05-10 | 31 | 3700 | 4279 | 4854 | 579 | 1154 |
| CA892 | Barguzin | M | 3+ | Intermediate | 2021 | 2021-08-30 | 2021-09-30 | 31 | 2022-04-07 | 2022-05-17 | 40 | 4024 | 4418 | 4834 | 394 | 810 |
| CA865 | Tunka | M | 2 | West | 2021 | 2021-08-18 | 2021-09-29 | 42 | 2022-04-20 | 2022-05-14 | 24 | 3362 | 4843 |  | 1481 |  |
| CA874 | Tunka | F | 2+ | West | 2021 | 2021-08-22 | 2021-09-30 | 39 | 2022-04-27 | 2022-05-19 | 22 | 3363 | 4444 | 4056 | 1081 | 693 |
| 27XK | Arhangai | M | 2 | West | 2023 | 2023-08-31 | 2023-10-08 | 38 | 2024-04-21 | 2024-05-15 | 24 | 3424 | 5384 | 4907 | 1960 | 1483 |
| 29AI | Arhangai | M | 2 | West | 2023 | 2023-08-29 | 2023-10-14 | 46 | 2024-04-19 | 2024-05-14 | 25 | 3021 | 4833 | 4455 | 1812 | 1434 |
| CA903 | Qinghai | F | 3+ | Highland | 2021 | 2021-09-11 | 2021-10-28 | 47 | 2022-03-27 | 2022-05-02 | 36 | 2032 | 2216 | 2536 | 184 | 504 |
| CA903 | Qinghai | F | 3+ | Highland | 2022 | 2022-09-11 | 2022-10-14 | 33 |  |  |  |  |  |  |  |  |
| CA905 | Qinghai | F | 2+ | Highland | 2021 | 2021-09-08 | 2021-10-01 | 23 | 2022-03-15 | 2022-05-01 | 47 | 1853 |  | 2691 |  | 838 |
| CA915 | Qinghai | F | 3+ | Highland | 2021 | 2021-09-11 | 2021-10-23 | 42 | 2022-04-13 | 2022-05-08 | 25 | 1686 | 2297 | 2851 | 611 | 1165 |
| CA917 | Qinghai | F | 3+ | Highland | 2021 | 2021-09-13 | 2021-10-14 | 31 | 2022-03-13 | 2022-04-30 | 48 | 1779 |  |  |  |  |
| CA921 | Qinghai | M | 3+ | Highland | 2021 | 2021-09-07 | 2021-10-19 | 42 | 2022-03-13 | 2022-04-20 | 38 | 1768 |  | 1899 |  | 131 |
| 29EH | Qinghai | M | 3+ | Highland | 2022 | 2022-09-12 | 2022-10-15 | 33 | 2023-03-25 | 2023-04-24 | 30 | 1991 |  |  |  |  |
| 29AN | Qinghai | F | 2+ | Highland | 2022 | 2022-09-09 | 2022-10-09 | 30 | 2023-03-22 | 2023-04-21 | 30 | 1952 | 2009 | 2482 | 57 | 530 |

a. The migration distance was calculated only when at least one stopover was detected in the SGAT analysis.

### 2. Migration phenology

Autumn migration occurred between late August to early October in the Russian and Mongolian populations, with an average duration of 38±7days; while spring migration occurred between early April to mid-May in these populations, with an average duration of 28±6 days. In the Qinghai population, autumn migration occurred between early September to late October, with an average duration of 35±7 days; and spring migration occurred between mid-March to early May, with an average duration of 36±8 days (**Figure 3**, **Table 2**).

**Figure 3.**
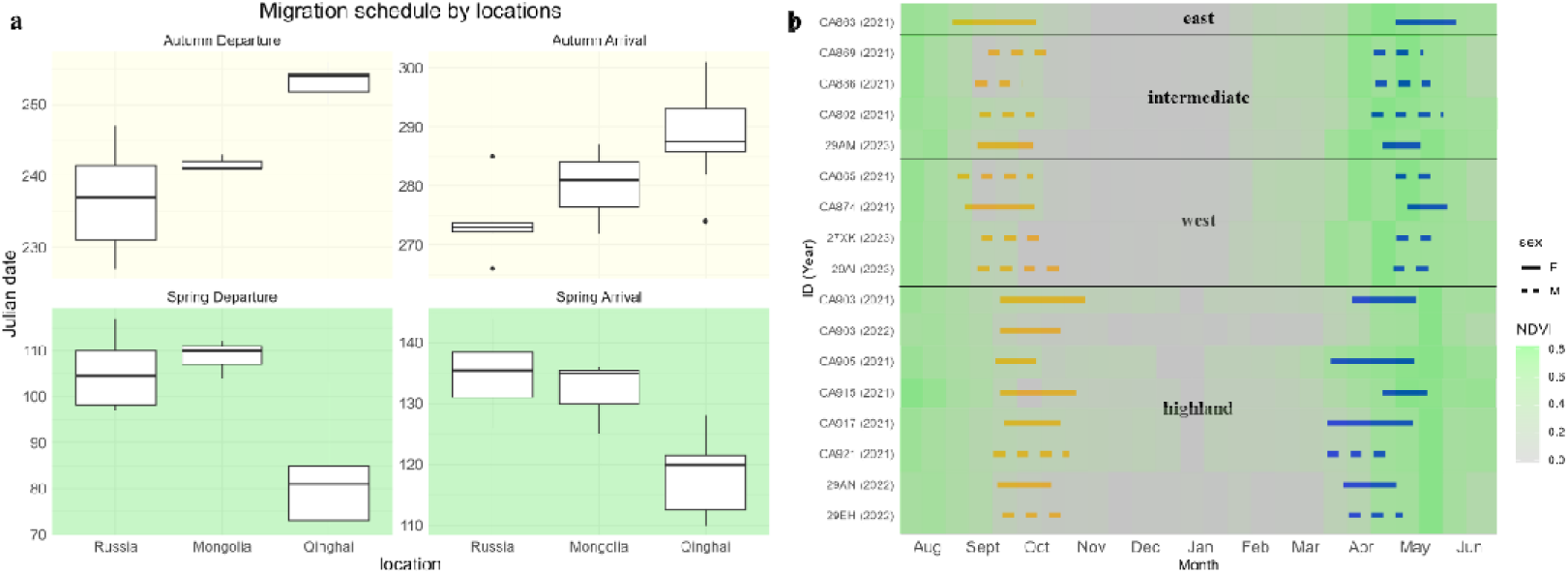
Migration phenology and its link to NDVI from breeding sites. **a.** Julian departure and arrival dates of autumn and spring migration, categorized by breeding locations. **b.** Migration schedules of all tracked individuals were grouped by four types of migration route (see Fig. 2). The bars represent the duration of the migration; autumn migration is in orange, and spring migration is in blue. Females are displayed with solid bars, whereas males are dashed bars. The background color represents the NDVI score (mean NDVI of 15 days, from 0 – 1) at the breeding site where the individual was caught. The NDVI annual datasets in each row were from the year when the individual was tracked.

### 3. Population structure and hybridization level inferred by WGS data

We acquired 18,837,382 SNPs after the extensive filtering after the variant calling step, and 594,881 SNPs were retained after the second round of filtering, with a genotyping rate at 0.995. One individual from the QTP (QTP_11) population and two from XIN (XIN_02, 03) were filtered out due to low data quality.

In the admixture analyses (**Figure 4**), the K=2 model (**Figure 4a**) held the lowest CV error value (0.4986±0.0003), where birds from the QTP (*przewalskii*) and from the westernmost KAZ+XIN (*maurus*) populations were categorized as two populations, respectively. All others were identified as their hybrids with different hybridization levels. The easternmost HLJ population (*stejnegeri*) has a genetic ancestry of 40 – 44% from the QTP population in this model. Individuals from the assumed Siberian contact zone in the Lake Baikal region, ILC, TUN, and BAR, carried 1 – 39.9% ancestry from the QTP population. The K=3 model (**Figure 4b**), which was our primary hypothesis according to the presumed taxonomy status (*maurus*, *stejnegeri*, *przewalskii*), had slightly higher CV error value (0.5190±0.0004) than K=1 (0.5074±0.0002) or 2 (0.4986±0.0003), but lower than K=4 (0.5670±0.0098) or 5 (0.6174 ± 0.0118) (**Figure S3**). In this model, QTP (*przewalskii*) and the westernmost KAZ+XIN (*maurus*) populations were again categorized as two populations, whereas the easternmost HLJ (*stejnegeri*) population was categorized as the third population. In the K=3 model, Eight of 17 individuals from the BAR population from the eastern Baikal region had complete *stejnegeri* genetic composition. The remaining birds carried a 2.5 – 40% genetic ancestry of *maurus*, while most remaining individuals were classified as *stejnegeri* based on their genetic composition. In the western Baikal ILC+TUN populations, no birds showed the complete *maurus* ancestry. For five of 20 individuals, the genetic ancestry of *maurus* was below 50% (4 – 41%), for another nine individuals it ranged between 50 – 75%, and for the remaining six individuals, ancestry of *maurus* was above 75% (79 – 97%). All but one individual (TUN_17) had a genetic ancestry of *stejnegeri* (12 – 96%).

**Figure 4.**
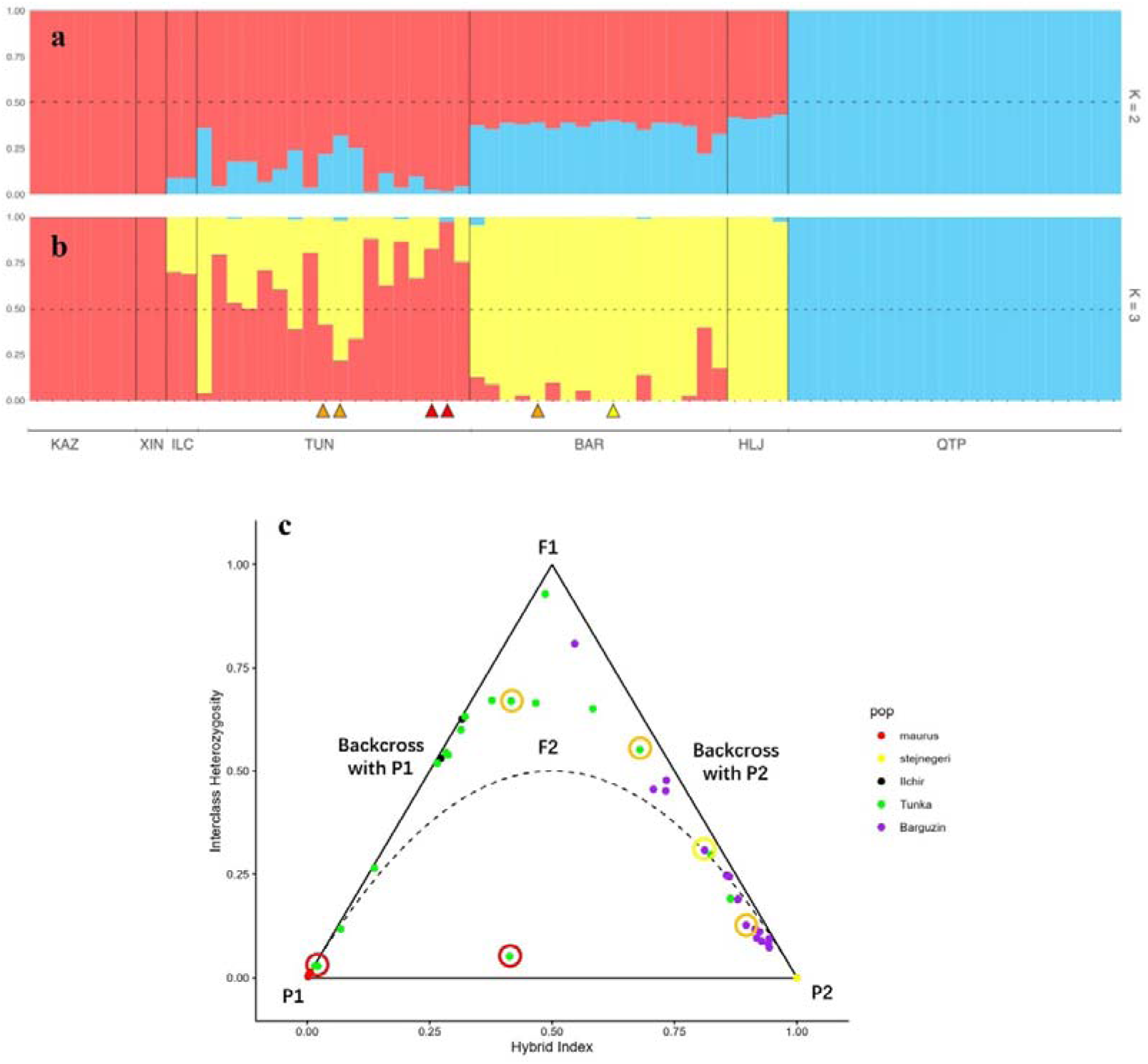
**a b**: The admixture plots (K = 2 and 3) of individuals from Kazakhstan, Xinjiang, Ilchir, Tunka, Barguzin, Heilongjiang and Qinghai, based on the whole-genome re-sequenced data after filtering. Each individual follows the same order in each plot. Black vertical solid lines separate individuals from different populations, with a reference to the population along the x-axis at the bottom of the plot. Black horizontal dotted lines indicate the 50% proportion of genetic ancestry. The yellow, orange and red triangles under the individual names and the corresponding bars refer to the individuals with tracking data, and the colors refer to the route groups: red represents the western route, orange represents the intermediate route, and yellow represents the eastern route. **c.** The triangle plot of the same group of individuals as included in the admixture plots, except for those from Qinghai. The plot is defined by solid lines: ylJ=lJ2x and ylJ=lJ−2xlJ+lJ2, and dotted curve ylJ=lJ2*(1lJ−lJx). The dotted curve represents the boundary below which individuals cannot occur, assuming Hardy-Weinberg Equilibrium. For parental groups, P1 were set as *maurus* using KAZ+XIN individuals, P2 as *stejnegeri* using HLJ individuals. The theoretical positions of different types of hybrids (F1, F2, backcross with P1, backcross with P2) were labeled on the triangle plot. Individuals with tracking data were circled, following the same color code as described in **a** and **b**.

To zoom into the contact zone between *maurus* and *stejnegeri,* the genetic composition of individuals was presented in a triangle plot. The parental group P1 was set as *maurus* using KAZ+XIN individuals, and the parental group P2 was set as *stejnegeri* using HLJ individuals (**Figure 4c**) The two individuals from Ilchir shared a pattern similar to a backcross with P1. Of the 17 individuals from Barguzin, 16 positioned between a pure P2 and a backcross with P2 on the plot; the remaining individual positioned close to an F1. Among the 18 Tunka individuals, one positioned close to an F1, three positioned close to a backcross with P2, nine positioned between a pure P1 and a backcross with P1 and four positioned close to be an F2; the one remaining individual could not be categorized due to an intermediate hybrid index combined with low interclass heterozygosity.

In the PCA plot (PC1∼PC2), the samples from the QTP population were clearly clustered and separated from the rest of the samples, on PC1. All other birds in the dataset were linearly distributed along a diagonal line between PC1 and PC2, ranging from the easternmost HLJ to the westernmost KAZ+XIN populations (**Figure 5a**). Consequently, both PC1 and PC2 showed a significant correlation with the breeding longitudes among this subset (**Figure 5b, c**).

**Figure 5.**
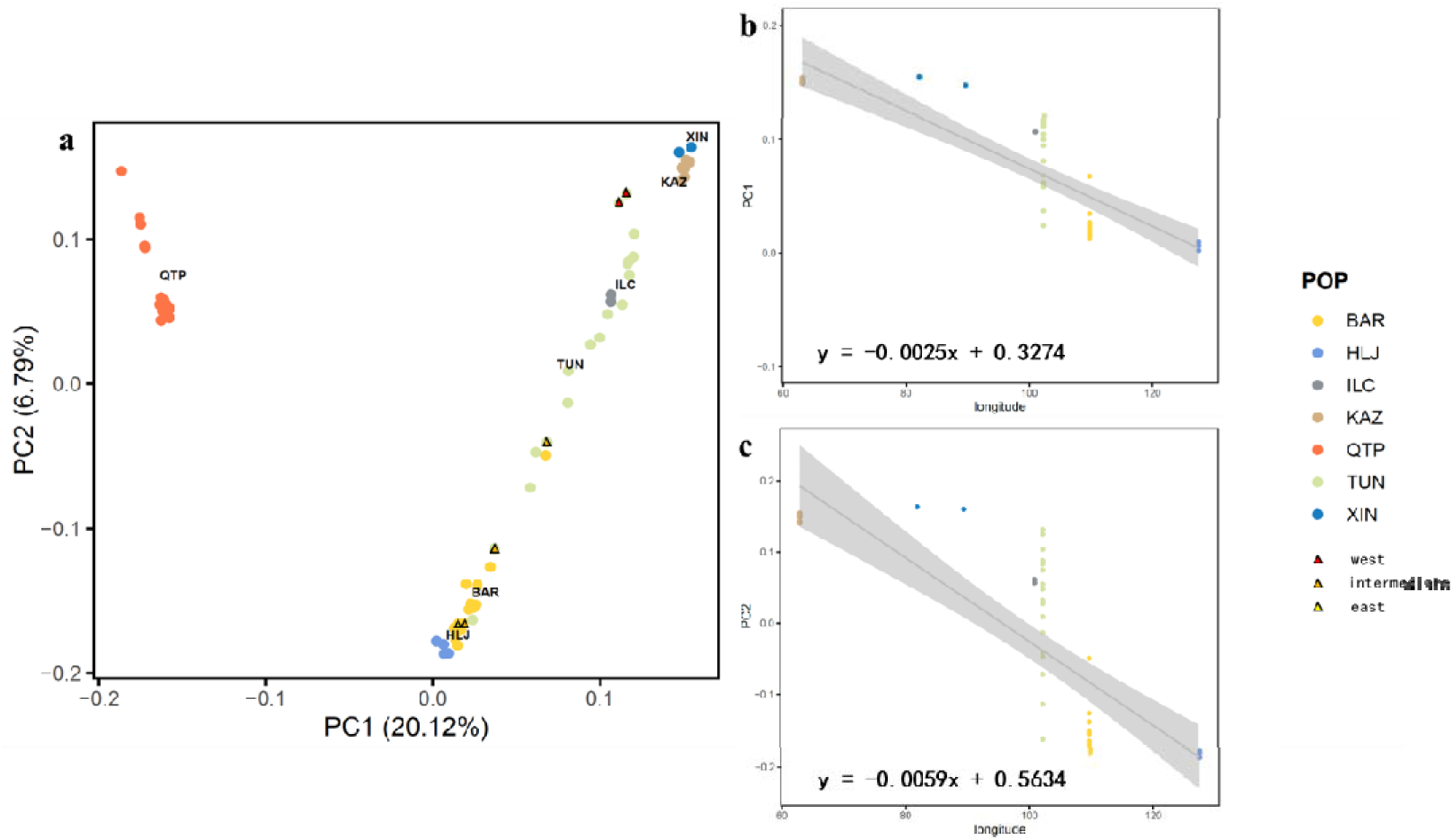
**a.** The PCA plot (PC1 ∼ PC2) of individuals from Kazakhstan, Xinjiang, Ilchir, Tunka, Barguzin, Heilongjiang and Qinghai, based on the whole-genome re-sequenced data after filtering. The percentage of variation explained by each PC is indicated in parentheses, and samples are coloured by population. For the six birds with both phenotypic and genotypic data from the contact zone, their dots were marked by triangles, with a color code referring to the route groups: red represents western route, orange represents intermediate route, and yellow represents eastern route. **b/c.** The relationship between PC1 (**b**) and PC2 (**c**) with breeding longitudes of all sampled individuals except for the QTP population. The colors of the dots refer to the same population color code as in **a**. The regression lines also show standard deviation, and equations are shown in the bottom-left corner in each plot.

The highest F_ST_ (0.13) was between the westernmost KAZ+XIN (*maurus*) and QTP (*przewalskii*) populations, whereas the easternmost HLJ (*stejnegeri*) and the east Baikal BAR populations shared the lowest F_ST_ at 0.00036. F_ST_ values for the remaining pairs were between 0.01 – 0.06 (**Table 3**). The easternmost HLJ (0.388) and westernmost KAZ+XIN (0.311) had higher observed heterozygosity values, whereas the three Baikal populations, BAR and TUN+ILC from the contact zone had lower heterozygosity (0.278 and 0.274, respectively), slightly higher than the level in the QTP population (0.254).

**Table 3.**
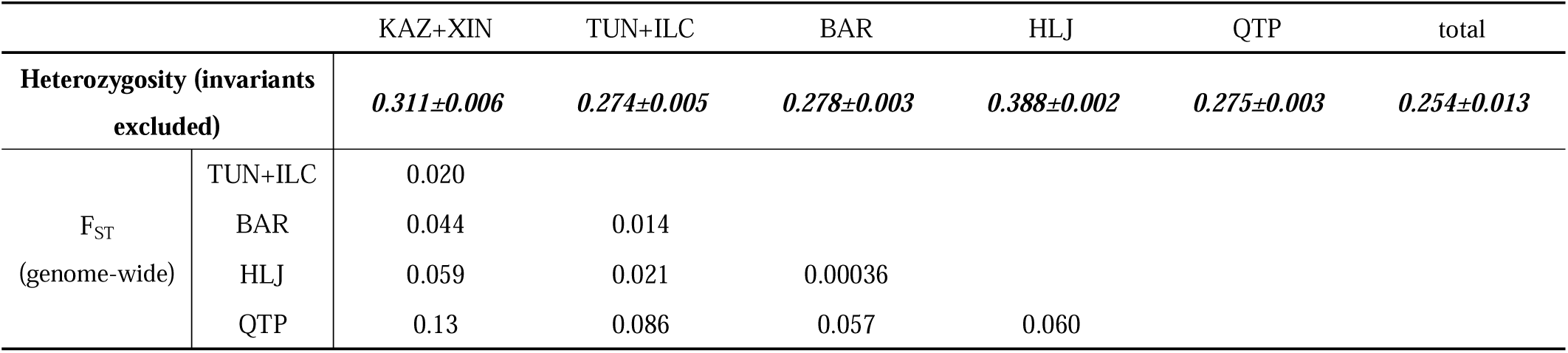
heterozygosity of each geographical breeding population, and genetic differentiation (F_ST_) between each pair of geographical breeding populations.

### 4. Relationship between migration route diversity and genetic variation

We labeled the route choices of the tracked individuals from the contact zone (Barguzin and Tunka) on the admixture plot, triangle plot (**Figure 4**) and PCA plot (**Figure 5**) using the same color code as in **Figure 2**. For the two individuals following the western route (western Baikal region TUN_16 and TUN_17), genetic components for *maurus* were 75% and 97% in the K=3 admixture model (**Figure 4**), and they were located very close to the westernmost KAZ+XIN populations’ cluster in the PCA plot (**Figure 5a**). They both had very low interclass heterozygosity level, despite the hybrid index of one of them being close to 0.5 (**Figure 5c**). The Barguzin individual following the eastern route was inferred to be a pure *stejnegeri* in the K=3 admixture model (**Figure 4**), and it clustered with the easternmost HLJ individuals in the PCA plot (**Figure 5a**). This individual positioned as a backcross type with *stejnegeri* in the triangle plot with an interclass heterozygosity near 0.25 (**Figure 5c**). The three individuals from either Barguzin or Tunka, which followed the intermediate routes showed 60 – 95% of genetic contributions of *stejnegeri* in the K=3 admixture model. These three individuals were all located between the western-routed and eastern-routed individuals in the PCA plot, but were closer to the HLJ cluster (**Figure 5a**). Two of the intermediate-routed individuals positioned in the range of F2 in the triangle plot, among which one was closer to a backcross with *maurus* while the other closer to a backcross with *stejnegeri*. Surprisingly, the third individual positioned very close to the parental *stejnegeri* type with lower interclass heterozygosity score than the east-route individual (**Figure 5c**).

## Discussion

Using individual tracking data, we confirmed the proposed existence of a migratory divide of stonechats in the contact zone between *maurus* and *stejnegeri* south of Lake Baikal (Irwin & Irwin, 2005) and showed that the divide extended to eastern Mongolia. Furthermore, we documented intermediate routes across the eastern part of the QTP, mostly by hybrids between *maurus* and *stejnegeri*. Moreover, we found that the *przewalskii* population breeding on the QTP migrated exclusively over the highlands during autumn migration, instead of starting the journey by leaving the QTP towards central China first, as recently described for the sympatric breeding population of Siberian Rubythroat (*Calliope calliope*) (Zhao et al., 2024).

Here, we presented an overall population genetics description of the Asian stonechats: the K=3 model in the admixture analysis, although not statistically optimal, supported a population structure referring to the pre-defined *maurus*, *stejnegeri* and *przewalskii* taxa. We also provide evidence for extensive hybridization in the contact zone between *maurus* and *stejnegeri* south of Lake Baikal. Because the genomic samples were all from adults that have completed two successful migration episodes back to the breeding site, we can infer that some hybrids were sufficiently viable to complete a full migration cycle and were engaged in reproduction. In other words, purifying selection is not complete against hybrids in the contact zone between *maurus* and *stejnegeri*, nor is there strong assortative mating by genotypes, indicating ongoing gene flow, opposing a speciation processes.

### Routes relative to geographical barriers in central Asia

In our study, we revealed the various migration route choices among stonechats in Asia, with seemingly different responses to the presumed geographical barrier, the Qinghai-Tibet Plateau. We hereby split the more detailed discussion for each route choice.

The route around the north-western side of QTP, i.e., the west route, has been documented in many bird species via tracking, e.g., the Demoiselle Crane (*Grus virgo*) (Galtbalt et al., 2022; Mi et al., 2022), and the Siberian Bluethroat (Strubbe et al., 2025). We showed that this route is also being used by stonechats. That this route encompasses a large detour of up to 59% compared to the bee-line distance (**Table 2**, **Figure 2**, **Figure S2**) may suggest certain benefits for the stonechats of taking it. A barrier-avoidance migration strategy may be an explanation for this pattern, where the QTP, along with other geographical barriers such as the Taklamakan Desert or the Tian Shan mountains, select against migration routes that directly cross these landscapes. Nonetheless, the documented western detour still passes unhospitable regions for the birds, such as the Tian Shan and Pamir mountains. In our data, we detected one stopover range in both seasons that was used by multiple birds (2/4 tracks in autumn and 2/4 tracks in spring), located around the Chirchiq River Valley, close to Tashkent, Uzbekistan (**Figure 2, Figure S1**). As an oasis along the route, it may function as a stopover refueling site and resting place where birds can wait for optimal weather and wind conditions before crossing unhospitable regions, or rest and refuel after barrier-crossing flights.

The route around the eastern side of QTP, i.e., the east route, involves a smaller detour of around 16% from the bee-line distance (**Table 2**); it was partially because the bird migrating along this route was found to select wintering locations further east, despite that we only had one track retrieved. This route resembles the previously documented migration path of Siberian Rubythroats (Zhao et al., 2024), and seemingly follows the mountain chain in central China. Lying between the Central Asia Flyway and the East Asia Flyway, central China may be frequently used by many species migrating on the east side of QTP (Heim et al., 2018, 2020; Kumar et al., 2020; Zhao et al., 2024).

Between the western and eastern routes, the intermediate routes used by many stonechats from the contact zone between *maurus* and *stejnegeri* passed the eastern side of the QTP and therefore crossed considerable elevations of over 3000m a.s.l.. We speculate that this behavior is facilitated by stonechat taxa being adapted to high elevations, as they are known to breed at high elevation ranges in Asia and Africa (Collar et al., 2020).

Parallel to the routes from stonechats breeding in Siberia and Mongolia, we document for the first time a “highland route” used by the population breeding on the QTP, corroborating our high-elevation adaptation hypothesis. The *przewalskii* stonechats are believed to breed both on QTP and along the mountains in central China, north up to Shanxi and Beijing, and south down to Yunnan (**Figure 1**); therefore, we had hypothesized that the QTP population would leave the highland, merging with the central Chinese populations during migration. However, none of the tracked individuals showed any signs of bypassing the QTP along its east side. Instead, they overlapped with the intermediate routes of southbound stonechats breeding in Siberia and Mongolia.

Stonechats breeding on the QTP, and northern breeders using intermediate routes, were all re-captured as breeders. This supports that for stonechats, the barrier effect of the QTP might not be as strong as predicted (Hawkes et al., 2011; Kumar et al., 2020). Interestingly, we did not document any migration trajectory crossing the western part of QTP, where the land is more barren (Liu et al., 2017) and the elevation is higher with more mountains over 5000m a.s.l.. This asymmetry may suggest that for stonechats, the actual barrier for optimal migration is limited to the western part of QTP. A greater barrier effect of the western QTP might also be linked to differences in vegetation and habitat availability: it consists more of alpine deserts, whereas more alpine meadows are found in the eastern part (Dong et al., 2020). Alternatively, the barrier effect from the Taklamakan Desert could play an important role for migratory landbirds, blocking the route between the western part of QTP and Siberia.

We would argue, however, that the ground elevation might not be the main contribution to the barrier effect, especially for species already breeding on high elevations, such as the *przewalskii* stonechats in our study. As mentioned in the introduction, the migration route along the eastern part of QTP may be shared by some other high-elevation breeding songbird species, e.g., the White-throated Bushchat, Blyth’s Pipit (*Anthus godlewskii*) and Rosy Pipit (Collar, 2020; Tyler et al., 2020); it was simply difficult to observe their actual migration passage due to low coverage of birders and low concentration effect at suitable stopovers on the QTP. Therefore, we propose that a careful revisit of the actual and region-specific barrier effect of the QTP and the surrounding landscapes should be undertaken, to understand how the landscapes in Asia have shaped the diversity of avian migration.

To expand the overview to the landbird migration in Central Asian Flyway, the migration patterns we described may be paralleled by species that are also suggested to have a Siberian migratory divide. For instance, Citrine Wagtail, White Wagtail (*Motacilla alba*) and Greenish Warbler (*Phylloscopus trochiloides*) were proposed to have a migratory divide (Irwin & Irwin, 2005), while they also share breeding populations on the Qinghai-Tibet Plateau. Migration patterns in these species may resemble the pattern of the stonechat taxa, where both highland routes and detoured routes around the QTP exist in parallel. We encourage further studies on different taxa to clarify the role of the QTP and surrounding barriers, to assess to what extent they have been shaping the migration diversity of landbirds in Asia.

### Population structure, hybrid zone property, and genetic perspective for the migration direction

From our genomic resequencing study, we infer frequent genetic exchange between *maurus* and *stejnegeri* in the contact zone. We demonstrated that most individuals from Ilchir and Tunka (19/20), and half of the individuals from Barguzin (9/17), had genetic ancestry of both *maurus* and *stejnegeri*, based on the admixture analysis (K=3) (**Figure 4**). Aside from *przewalskii* stonechats, there was a clear linear relationship between both PC1 and PC2 with the breeding longitude, from pure *maurus* (Kazakhstan and Xinjiang, China) to pure *stejnegeri* (Heilongjiang, China) populations (**Figure 5b, c**). Low F_ST_ along these Siberian populations also indicated low fixation level of population genetic features (**Table 3**). We therefore shift the terminology from “contact zone” to “hybrid zone”.

The hybridization at the hybrid zone between *maurus* and *stejnegeri* might be extensive. In the admixture plot (K=3), the high frequency of individuals with around 25% and 75% genetic contributions of *maurus* or *stejnegeri* supported the existence of F2 hybrids and backcross individuals (**Figure 4b**). In the triangle plot (**Figure 4c**), a substantial number of individuals was categorized as F2 and backcrosses in the hybrid zone, also suggesting that there is prolonged and ongoing genetic mixing over many generations.

These indications for continuous admixture may have contributed to the lower-than-expected number of predicted populations by the STRUCTURE model (Lawson et al, 2018). Additionally, if the sampled populations were actually derived from an ancestral population or lineage that was *not* sampled in our study, a so-called “ghost” population, this could also lead to an under-estimation of the number of populations (Lawson et al, 2018). To resolve the potential existence of a ghost population, we would require genomic sequences of a more extensive set of stonechat species, so that ancestry can be inferred in a more global phylogenetic context.

As a note from the field, five out of six Tunka and Barguzin re-captured individuals were actively breeding with nestlings when re-captured in 2022. CA874 (TUN_16, west route, female, arriving May 19th) and CA886 (TUN_07, intermediate route, male, arriving May 10th) were even forming a breeding pair in 2022, despite the fact that they were paired with other mates in 2021. Based on these observations, we would propose that there is no strong assortative mating preventing the hybridization between different migration phenotypes or genotypes. However, as there were several individuals in the triangle plot positioning below the dotted curve, we should assume that the hybridization between *maurus* and *stejnegeri* violated the expectation from Hardy-Weinburg Equilibrium (Wiens et al., 2025); natural selection or drifts may be expected among the selected 647 diagnostic loci for the triangle plot. Additionally, the sample size and strong filtering may provide an incomplete overview of hybridization occurrences, for example, when the diagnostic alternate alleles are not fixed in the parental groups, or due to technical batching effects.

Our results also enabled a better description of the range of the hybrid zone, which previous studies using haplotyped mitochondrial DNA data could not show (Illera et al., 2008; Opaev et al., 2018; Zink et al., 2009). The relatively uniformed “*stejnegeri*-dominant” genotype of stonechats at Barguzin, east of Lake Baikal (**Figure 4**), and its low F_ST_ score compared with the easternmost Heilongjiang population, suggest that Barguzin is close to the border of the hybrid zone where the eastern genotype prevails. On the western side, our study sites in Tunka valley and at Ilchir Lake are still within the hybrid zone, albeit with diminished contributions of *stejnegeri*. Local birds in this zone phenotypically resemble pure *maurus* (M. Hellström, personal observation). Therefore, the western border of the hybrid zone must be further extended to the west. Despite the lack of whole-genome re-sequencing data from the Mongolian individuals (Khurkh and Arhangai), the presence of an individual migrating along an intermediate route at Khurkh suggests that the hybrid zone between *maurus* to *stejnegeri* could potentially extend into eastern Mongolia. More sequencing data in the future would help to clarify the range of the hybrid zone. Overall, our findings are in disagreement with the argumentation by Opaev et al., 2018 that the transition between *maurus* to *stejnegeri* in Transbaikalia is abrupt. Due to the massive representation of intermediate genotypes, the identification by plumage and songs in the contact zone may not be an effective and accurate measure for species identification.

In our results, an association between the intermediate-route phenotype and hybrid genotype was suggested: the two intermediate-route individuals from Tunka carried a “*stejnegeri*-dominated” hybrid genotype in the admixture model (**Figure 4b**) and both were categorized as F2 or backcross hybrids (**Figure 4c**); whereas the intermediate-route individual from Barguzin carried a full *stejnegeri* ancestry in the admixture model (**Figure 4b**) and positioned very close to the *stejnegeri* parental type in the triangle plot (**Figure 4c**).

Since the initial proposal of the hypothesis regarding hybrids taking intermediate routes from a migratory divide (Helbig, 1991), investigations have been carried out in several systems (Delmore & Irwin, 2014; Delmore et al., 2020a; Zhao et al., 2020; Sokolovskis et al., 2023; Zając et al., 2024). Despite various genomic signals detected to be associated with the intermediate route behavior, there hasn’t been any particular gene identified to be responsible to the migration direction trait. To understand the genetic basis of migration direction, future studies in the Asian stonechat system will require the collection of more migration tracking and genomic resequencing data. Furthermore, a high-resolution annotated reference genome of stonechats should also be aimed for, in order to facilitate the annotations of candidate genes (Weissensteiner et al., 2025).

In addition to exploring the genetic mechanism responsible for the phenotypic trait variation, the Asian stonechat system further provides opportunities to study the relationship between migration patterns and historical expansion history. Irwin et al. (2005) and Irwin & Irwin (2005) demonstrated the Greenish Warbler and Two-barred Warbler (*Phylloscopus plumbeitarsus*) from both a ringed speciation and a migratory divide perspective. Though not directly inferred, the formation of the migratory divide between the Greenish and Two-barred Warblers could be assumed as a result of the ringed speciation. A similar investigation was conducted in the Willow Warbler system, where two contact zones between the subspecies *P*. *t*. *trochilus* and *acredula* were assumed to be associated with two migratory divides (Bensch et al., 2009). In both cases, the secondary contact between two closely-related taxa with different migration phenotypes were assumed to come with a post-zygotic reproductive isolation induced by the suboptimal intermediate route.

However, the Asian stonechat system seems not to follow either of the patterns in the warblers’ systems discussed above. Firstly, different to the Greenish Warbler, whose QTP population was more closely related to the western Siberian breeding population, the *przewalskii* stonechat seemed to share a closer relationship with *stejnegeri*, the eastern Siberian breeding population (**Table 3**). This suggests a different expansion history of breeding range between two taxa. Inclusion of the subspecies *indicus* (**Figure 1**) in the genetic study is needed to re-construct the historical expansion of stonechats in Asia. Secondly, the seemingly viable intermediate route phenotype, extensive hybridization, and low F_ST_ between *maurus* and *stejnegeri* also indicate a complex scenario, which may not be perfectly explained by a ringed speciation scenario, that *maurus* and *stejnegeri* have secondary contact where the migratory divide lies.

An alternative scenario may apply that the migratory divide can be formed *in-situ*, i.e., the migration phenotype shifted abruptly during the historical population expansion. In the best supported K=2 admixture model, the Ilchir to Heilongjiang populations were all inferred to be hybrids between *maurus* and *przewalskii* (**Figure 1**, **Figure 4a**); the high correlation between PC1 and PC2 with breeding longitudes between *maurus* and *stejnegeri* (**Figure 5b, c**) can also be explained as a single continuous population. Even though we cannot yet differentiate which scenario applies to the stonechat migratory divide, we propose that an *in-situ* occurrence of a migratory divide is a possible hypothesis. In fact, new migration directions can emerge in bird populations in a very short range of time, e.g., in the Richard Pipits (*Anthus richardi*), Yellow-browed Warblers (*Phylloscopus inornatus*) and Eurasian Blackcaps (Dufour et al., 2021, 2022; Helbig et al., 1994). We believe that future studies of demographic history in Asian stonechats, including more sampling sites, e.g., from the *indicus* range will help us to further untangle the relationship between migration phenotypes and biogeographical patterns. Studies on other species with a presumed Siberian migratory divide, where data is largely lacking, are also important to help us understand the evolution of landbird migration in Asia.

### Conclusions

Our study revealed the diversified migration phenotype, and its associated genotypic variations, among stonechats in Asia. The differential interaction between route choices and the geographical barriers may have resulted in genomic differentiation, but hybridization and survival of intermediate migration phenotypes dampened a speciation process and may provide sources for future evolution.

Our study complements investigations on the genetic basis of migration in a novel system, in parallel to established systems from Europe and North America (Caballero-López & Bensch, 2024; Justen & Delmore, 2022; Liedvogel et al., 2011), as well as Central Asia (Turbek et al., 2022). The different expansion histories and the interaction process between barrier landscapes and evolution in different continents/flyways could provide multiple systems to assess whether the evolution of migration phenotypes converged or not. Since the evolutionary history of migration may vary among systems, it is important to involve diverse systems, including the so far under-represented Asian flyways.

## Acknowledgement

We thank Hongyu Zhao and Suqin Hu for their financial support on the fieldwork conduction and whole genome re-sequencing. We thank Aleksey Bezrukov, Olga Mikhaleva, Oksana Fomina, Elmira Zainagutdinova, Aisha Anisimova, Otgonjargal Surakhbayar, Andrey Anisimov, and Danila Bezrukov for their assistance during the Russian and Mongolian fieldwork. We thank the following field assistants and volunteers that helped with the fieldwork in Qinghai, China: Xinyue Chen, Zhiying Du, Yangfan Feng, Chang Gao, Toto Liang, Boye Liu, Danmeng Liu, Jing Luo, Qiujin Luo, Chen Lv, Bread, Xiaotong Ren, Theo, Tony, Liufeng Wang, Liuyang Wang, Shan Wang, Xiaodan Wang, Yiqing Wang, Yushan Wang, Wanyi Wei, Yuning Wei, Shiyang Wu, Yinan Wu, Yaxi Xie, Chao Xing, Nuoyan Xu, Wanghong Yang, Teng Ye, Lang Yu, Yifan Yue, Ying Zeng, Baoling Zhang, Dalei Zhang, Shen Zhang, Xiangyu Zhao, Yuetao Zhong, Anqiang Zhou, Larry Zhuang. We thank Yayue Gao, Yu Zhang, Ms. Ding, Ms. Miao and Haosheng Xu for their coordination on the fieldwork permits. We thank Mr. and Mrs. Ma from Haoqing hotel for their hospitality during our fieldwork. We thank Chinese Bird Banding Association and Lixia Chen for the provision of metal rings. We thank Zdeněk Moudrý for the supply of perch traps. We thank Ecotone.pl for their supply of color rings. We thank Tiancheng Gao, Lu Dong and Xi Huang for their help for sample transportation and deposition in China during Covid-19. We thank Steffen Hahn for his assistance on SOI logger consultation, and Neringa Znakovaité for her assistance on SOI logger data extraction. We thank Martins Briedis, Joanna Wong, and Kristaps Sokolovskis for their important help during the geolocator analysis. We thank Swiss Ornithology Institute and Migrate Technology ltd. for their logger production. We thank Per Alström and Yang Liu for their inputs during the project conceptualization phase. We thank Jun Ishigohoka and Corinna Langebrake for important advice and assistance on bioinformatic analyses. We thank Anna Rensink for the help in the lab. We thank Peter de Knijff for his suggestion on barcoding in *Saxicola* taxa, and his insightful comments on the Siberian contact zone of stonechats. We thank Shuangqi Liu for the migrants’ information from Motuo county, and the constant discussions. We thank Wechat group “Erzicun” for commenting on the figures. We thank the constant academic discussion with Hongkai Zhang, Chuan Jiang, Xiaotong Ren, Jacob Roved, Shujie Liang and Xi’er Chen.

## Authors’ contribution

TZ, BH, BW, MH, FL, GS, ML, KR, SB and WH together conceptualized the study and initiated the project. TZ, DZ, GS, GZ, ZL, WC, SJ, D, YZ, XW, SL, SC, AL, ZL and GW contributed at least two years of full-season fieldwork in Qinghai, China. YA and VA organized and conducted the fieldwork in Russia. NB, BD, YA, VA and WC organized and conducted the fieldwork in Mongolia. TZ and WH conducted the geolocator analyses. FL, GS, ML, AB, BH provided and organized tissue or specimen samples from Kazakhstan, Xinjiang and Heilongjiang; GS, XJ, DZ, JR and TZ conducted the labwork and shipment for whole-genome re-sequencing. MW, GL, GW, KR, CH and TZ prepared the materials and conducted the bioinformatic analyses. TZ wrote the manuscript with the contribution and revision by all other co-authors.

## Availability of data and materials

The geolocator data, both the raw data and processed geolocation data, are archived in a Movebank repository: https://doi.org/10.5441/001/1.705. All the genomic re-sequencing data will be upload to Genbank soon and can be requested on demand.

