## Supplementary figures and images for "Migration patterns and hybridization within the Asian stonechat complex in response to a major geographical barrier"

### Figure S1

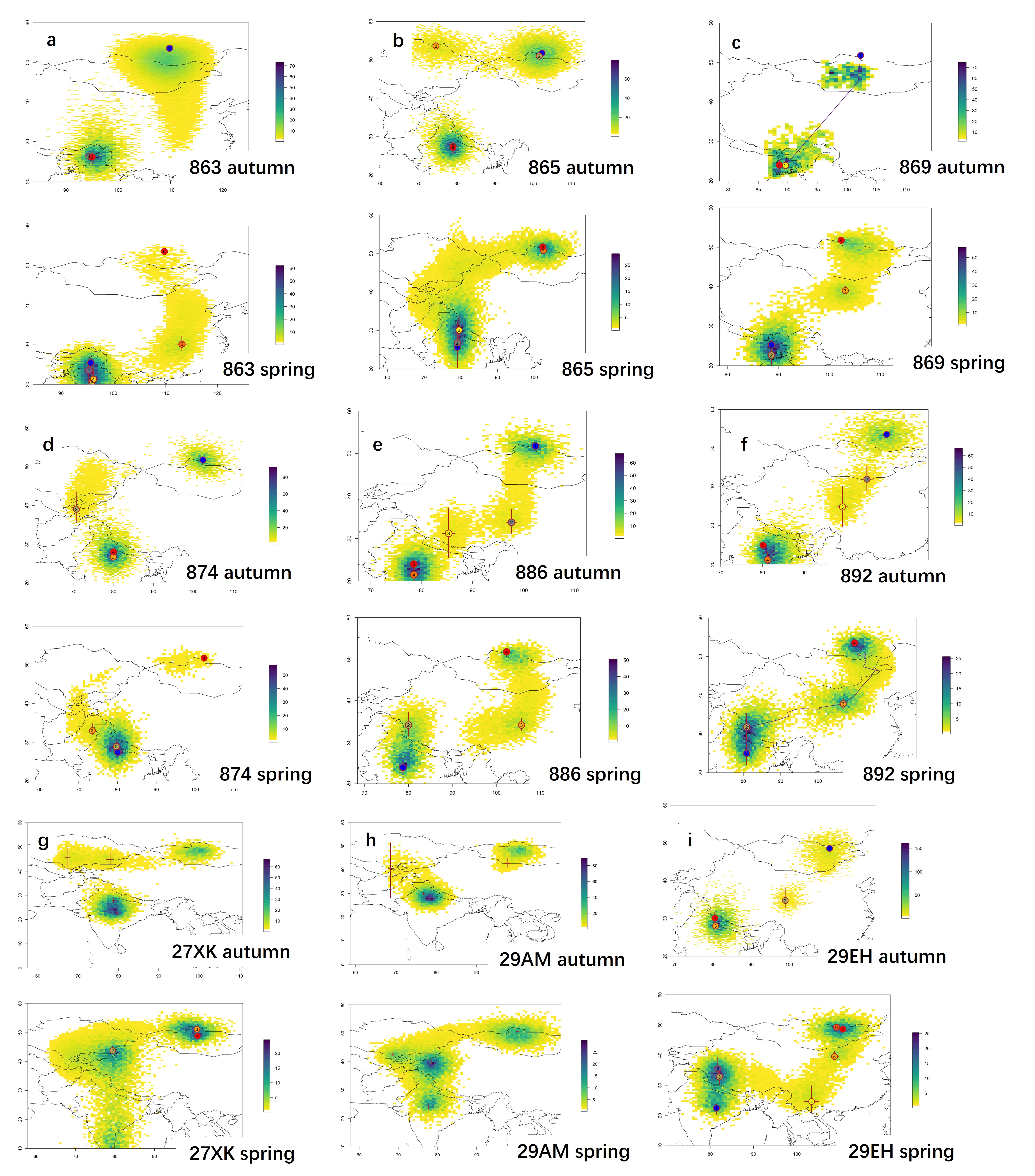

### Figure S2

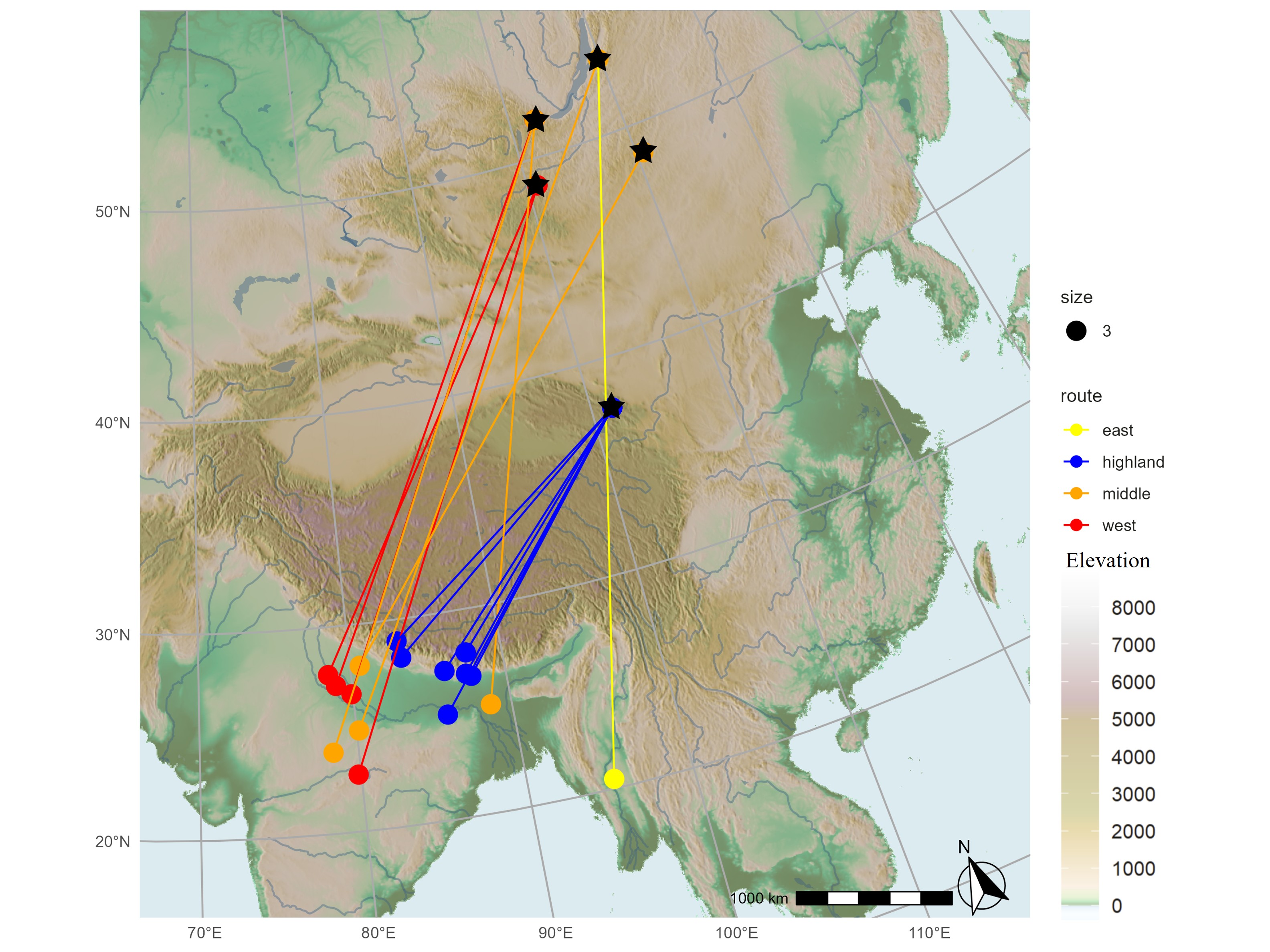

### Figure S3

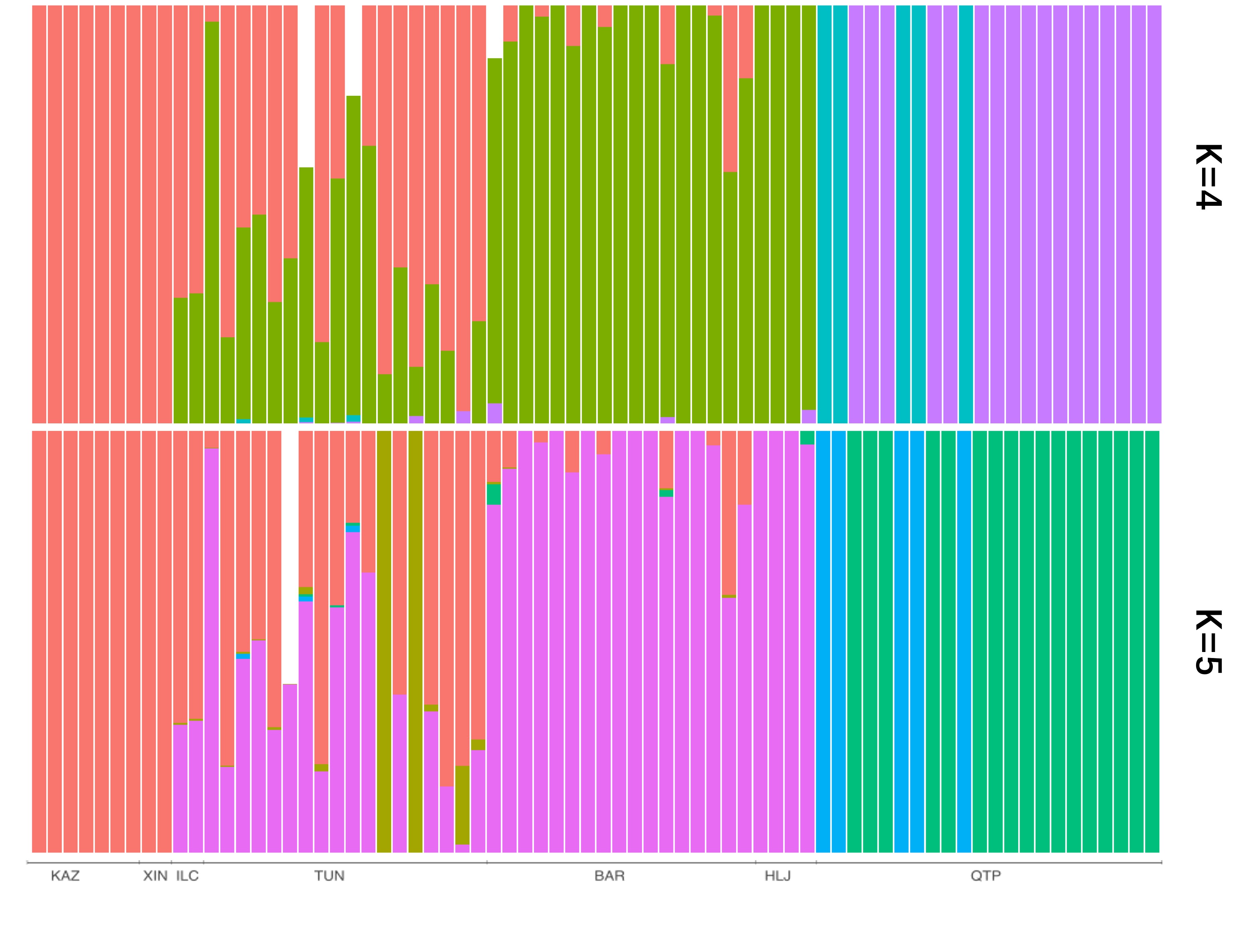
