## Supplementary material for "Migration patterns and hybridization within the Asian stonechat complex in response to a major geographical barrier"

Supplementary materials

- Supplementary methods
- Supplementary Figure 1
- Supplementary Figure 2
- Supplementary Figure 3
- Supplementary Table 1
- Supplementary Table 2

**Supplementary methods**

#Geolocator data analysis protocol

We first log-transformed the light-level data using the preprocessLight() function in the TwGeo package to annotate and edit the twilights. For calibration, we used the thresholdCalibration() function with a defined period before departure from the breeding site to acquire the zenith (summer), and using the findHEZenith() function using a winter period to acquire an alternative zenith (winter). All the calibration settings are enclosed in Table S1.

When conducting the SGAT analysis, we first split each dataset into autumn and spring on January 1^st^ or February 1^st^ in the corresponding years to run the SGAT analyses separately. The dataset CA903 had 1.5 years of data, so we split it into two autumn and one spring dataset. For all the Russian and Mongolian birds we used the summer zeniths to calibrate the data, as the summer and winter zeniths did not differ more than 0.5° in these datasets. For the Qinghai birds, since the summer and winter zeniths differed above threshold levels (>1°), we either used the winter zeniths for both seasonal datasets, or used the summer zeniths for the autumn data and winter zeniths for the spring data. The winter zeniths gave more precise estimations for wintering locations; thus, we used the inferred winter locations from the spring datasets to illustrate the migration trajectories. We also optimized the settings of quantiles (0.9 – 0.97), threshold of duration of days (5 or 10) in the function changeLight(), as well as the distance threshold (500 km or 1000 km) in the function mergeSites() for each individual to enable the performance of the group movement model in the SGAT analysis (**Table S1**). For the movement model, we assumed a Gamma distribution of speed with k=2.2 and λ=0.08, which allowed the movement to be steadily continuous and fewer stationary sites to be detected. With these settings we cannot detect all short stopovers; but the detected stopovers are more robust. For each dataset from Russia and Mongolia, we tried to adjust the settings of e.g., quantiles in the changeLight() function to detect at least one stopover during each migration season to determine whether the bird had detoured from the beeline between breeding and wintering site. To quantify the difference between the migration distance and the bee-line distance between breeding and wintering sites, as well to compromise to the low resolution of migration routes inferred from geolocator data, we used the longer distance from either autumn or spring migration for each individual to represent its migration distance. We only considered the routes with at least one detected stopover for the migration distance calculation.

We inferred the migration phenology pattern using the functions changeLight() (quantile = 0.8, day = 2) and mergeSites() (distance = 500 km). We only extracted the start and end date of autumn and spring migration, respectively.

#Pre-processing of genomic data

The variant-calling pipeline comprises the following steps: 1. Adaptor-trimming, quality-based filtering, and error-correction of reads with *fastp* (Chen et al., 2018; Chen, 2023), 2. Mapping of reads to the stonechat reference genome (accession GCA_900205225.1) with *bwa-mem2* (Vasimuddin et al., 2019), 3. Duplicate removal, alignment file processing with *samtools* (Danecek et al., 2021), 4. Variant calling with *bcftools mpileup* and *bcftools call* (Danecek et al., 2021), and 5. Extensive filtering of the produced variants (keep only bi-allelic SNPs with depth between 5 – 600, exclude genotypes with maf < 0.0000001 and hwe < 1e-50 to minimize calling bias, GQ value > 30; exclude loci with genotyping missing rate > 10%, exclude individuals with missing rate > 30%).

**Supplementary tables (captions)**

**Table S1** Light-level data analysis settings for each individual data set.

**Table S2** Genomic re-sequencing sample list and technical information

**Supplementary figures (captions)**

**Figure S1** Inferred migration trajectory results from SGAT summary analyses for all birds tracked from Russia and Mongolia. Each individual has one plot for autumn and another for spring. In each plot, the detected stationary sites (including breeding site) were circled in different colors, with a number sequence indicating the temporal sequences of the occurrence for these stationary sites. The distribution likelihood of the trajectory is shown in each plot: the darker color, the more likely the birds occurred. This figure provides more direct evidence of the seasonal usage of routes.

**Figure S2** The Great-circle lines of all tracked individuals from their breeding sites to the wintering sites. The breeding sites were marked with black stars, whereas the wintering sites were marked with circles. The color of the lines and circles followed the same color codes as Figure 2: yellow represents east route, orange represents intermediate route, red represents west route, and blue represents highland routes.

**Figure S3** The K=4 and K=5 model outputs from the admixture analyses. Different colors represent different assignments of population groups.
